# RNA polymerases IV and V influence the 3’ boundaries of Polymerase II transcription units in *Arabidopsis*

**DOI:** 10.1101/108902

**Authors:** Anastasia McKinlay, Ram Podicheti, Jered M. Wendte, Ross Cocklin, Douglas B. Rusch

## Abstract

Nuclear multisubunit RNA polymerases IV and V (Pol IV and Pol V) evolved in plants as specialized forms of Pol II. Their functions are best understood in the context of RNA-directed DNA methylation (RdDM), a process in which Pol IV-dependent 24 nt siRNAs direct the *de novo* cytosine methylation of regions transcribed by Pol V. Pol V has additional functions, independent of Pol IV and 24 nt siRNA biogenesis, in maintaining the repression of transposons and genomic repeats whose silencing depends on maintenance cytosine methylation. Here we report that Pol IV and Pol V play unexpected roles in defining the 3’ boundaries of Pol II transcription units. Nuclear run-on assays reveal that in the absence of Pol IV or Pol V, Pol II occupancy downstream of poly A sites increases for approximately 12% of protein-coding genes. This effect is most pronounced for convergently transcribed gene pairs. Although Pols IV and V are detected near transcript ends of the affected Pol II – transcribed genes, their role in limiting Pol II read-through is independent of siRNA biogenesis or cytosine methylation. We speculate that Pols IV and V (and/or their associated factors) play roles in Pol II transcription termination by influencing polymerase bypass or release at collision sites for convergent genes.

## Introduction

Nuclear multisubunit RNA polymerases IV and V (Pol IV and Pol V) play distinct roles in the transcriptional silencing of genes and transposons via RNA-directed DNA methylation (RdDM)^1^. Pol IV is thought to initiate the major RdDM pathway, transcribing DNA to generate relatively short RNA transcripts that are then copied into double-stranded RNAs by RNA-DEPENDENT RNA POLYMERASE 2 (RDR2)^2–9^. Resulting double-stranded RNAs are diced by DICER-LIKE 3 (DCL3) into 24 nt siRNAs that are loaded into ARGONAUTE 4 (AGO4), or a related AGO protein^10–13^. In parallel, Pol V generates transcripts and recruits AGO4-siRNA complexes to RdDM loci via its C-terminal AGO4 hook motifs^14–16^. As Pol V elongates, AGO4-siRNA interaction with either complementary bases in lncRNA or DNA guides *de novo* cytosine methyltransferase, DRM2, to bring about repressive cytosine methylation (in CG, CHG or CHH contexts)^2,16–19^. There is evidence that Pol V has functions independent of RdDM, helping maintain repression of transposons and genomic repeats whose silencing depends on maintenance methylation by MET1 and the chromatin remodeling ATPase, DDM1^20–22^. First, it has been reported that heterochromatin organization and silencing of pericentromeric transcription is affected in *pol V* mutant plants independently of the 24-nt siRNA biogenesis pathway^20^. Also, Pol V has been implicated in at least one alternative TE silencing pathway, the RDR6-dependent DNA methylation pathway (RDR6-RdDM)^12,21–24^. In this Pol IV- and 24-nt siRNA-independent pathway, Pol II-derived RNAs from TE regions are processed into 21- and 22-nt siRNAs in a RDR6, DCL2, and DCL4-dependent manner, loaded into AGO6 and targeted to Pol V scaffold transcripts^21,22,24^. Thus, Pol V can function as a downstream component in both 24-nt siRNA-dependent (canonical RdDM) and 21- and 22-nt siRNA-dependent (RDR6-RdDM) TE silencing pathways.

Steady-state levels of mRNAs transcribed by Pol II in *Arabidopsis thaliana* are minimally affected in *pol IV* or *pol V* mutant plants^25^. However, several observations have suggested that Pol IV and V function in proximity to Pol II transcription units. At long retrotransposons, 24-nt siRNA biogenesis and RdDM occur primarily at the ends of the elements, where promoter sequences recognized by the Pol II transcription machinery are located^21,26^. Moreover, chromatin immunoprecipitation studies have shown that approximately 30% of Pol V occupied positions are proximal to the promoters of Pol II - transcribed genes^17^. Likewise, 21-24 nt sRNAs are enriched within 1 kb of either the 5’ or 3’ ends of transcription units for approximately 20% of protein-coding genes, with studies suggesting that sRNA abundance correlates with mRNA abundance at these genes^27,28^. However, unlike Pol V ChIP peaks that correspond to transposons, Pol V peaks that overlap protein-coding genes typically do not correlate with 24-nt siRNAs or CHH DNA methylation^17^, which are hallmarks of RdDM, suggesting the possibility of an alternative function of Pol V in regulation of protein-coding genes.

RNA Pol II transcription termination in eukaryotes is a complex process. Termination by Pol II occurs stochastically within a region of several hundred bases downstream of the polyA site and is functionally connected to the cleavage and polyadenylation of nascent transcript 3’ ends^29–32^. Transcription pause sites, binding sites for protein termination factors, the presence of RNA-DNA hybrids, or chromatin modifications within the termination region can affect termination efficiency^33–35^.

We conducted nuclear run-on assays combined with high-throughput sequencing (NRO-Seq)^36^ to identify genomic regions where nascent RNAs are synthesized, comparing wild-type plants to *pol IV* or *pol V* mutant plants, similar to an analogous study conducted in maize^37^. Interestingly, as in maize, we detect increased RNA Pol II abundance downstream from the cleavage and polyadenylation sites of protein coding genes in *pol IV* or *pol V* null mutants. By contrast, nascent transcript abundance in the region from the initiation site to the PolyA site is unaffected, as are steady-state mRNA levels. This suggests that 3’ end processing of pre-mRNAs is not affected in *pol IV* or *pol V* mutants, but termination and Pol II release is impaired or delayed. Convergently-transcribed protein coding genes whose 3’ regions overlap show the greatest increase in nascent RNA signals downstream of PolyA addition sites when Pol IV or Pol V are mutated. Those gene pairs also have a higher tendency to overlap with sites of Pol IV, Pol V or AGO4 occupancy, as measured by ChIP. However, these regions are not characterized by abundant 24-nt siRNAs or CHH methylation, suggesting that Pol II transcription termination does not require RdDM modification within termination zones. Collectively, our results suggest a methylation independent role for Pols IV and V in shaping the boundaries of Pol II transcription units.

## Results

### Nascent transcript levels increases downstream of mRNA polyA addition sites in *pol IV* or *pol V* mutants

Nascent RNA transcripts produced by template-engaged RNA polymerases in purified nuclei were generated by nuclear run-on (NRO) in the presence of biotin-UTP, allowing capture of the labeled RNAs using streptavidin beads. Quantitative PCR (qPCR) assays, comparing reactions conducted using biotin-UTP versus unmodified UTP, were then used to calculate the fold enrichment achieved for biotinylated RNAs versus background for a set of highly-expressed *Arabidopsis thaliana* genes (Suppl. Figure 1). This analysis allowed us to estimate that nascent biotinylated RNA accounts for more than 99.7% of the RNA captured on the streptavidin beads, with background binding, attributable to unlabeled RNAs, accounting for the remaining 0.3%. We performed nuclear run-on using nuclei of wild-type plants as well as *pol IV* and *pol V* mutant plants and constructed sequencing libraries for resulting RNAs (NRO-Seq library), as well as for purified total nuclear RNAs (RNA-Seq library), thus allowing comparisons between nascent and steady-state RNA levels (Suppl. Figure 2). To examine distribution of RNA polymerase II activity we focus our analysis on 1 kb regions surrounding RefSeq transcription start sites (TSSs) or transcription end sites (TESs) for 27859 protein-coding genes (Figure 1). Both nascent RNA (Figure 1A, **in black**) and steady-state RNA (Figure 1B, **in black**) profiles show a sharp increase in RNA sequence reads starting at the TSS, followed by a plateau that begins ∼200 bps downstream. This pattern contrasts with nascent RNA profiles reported for yeast and other eukaryotic organisms, in which peaks are often observed downstream of the TSS, reflecting promoter-proximal pausing of Pol II^36,38–42^. The absence of a promoter-proximal peak in the nascent RNA profile in *Arabidopsis* suggests that promoter-proximal pausing is not a general feature of *Arabidopsis* genes transcribed by Pol II. Similarly, no evidence of promoter-proximal pausing was observed in maize NRO data^37^.

**Figure 1:**
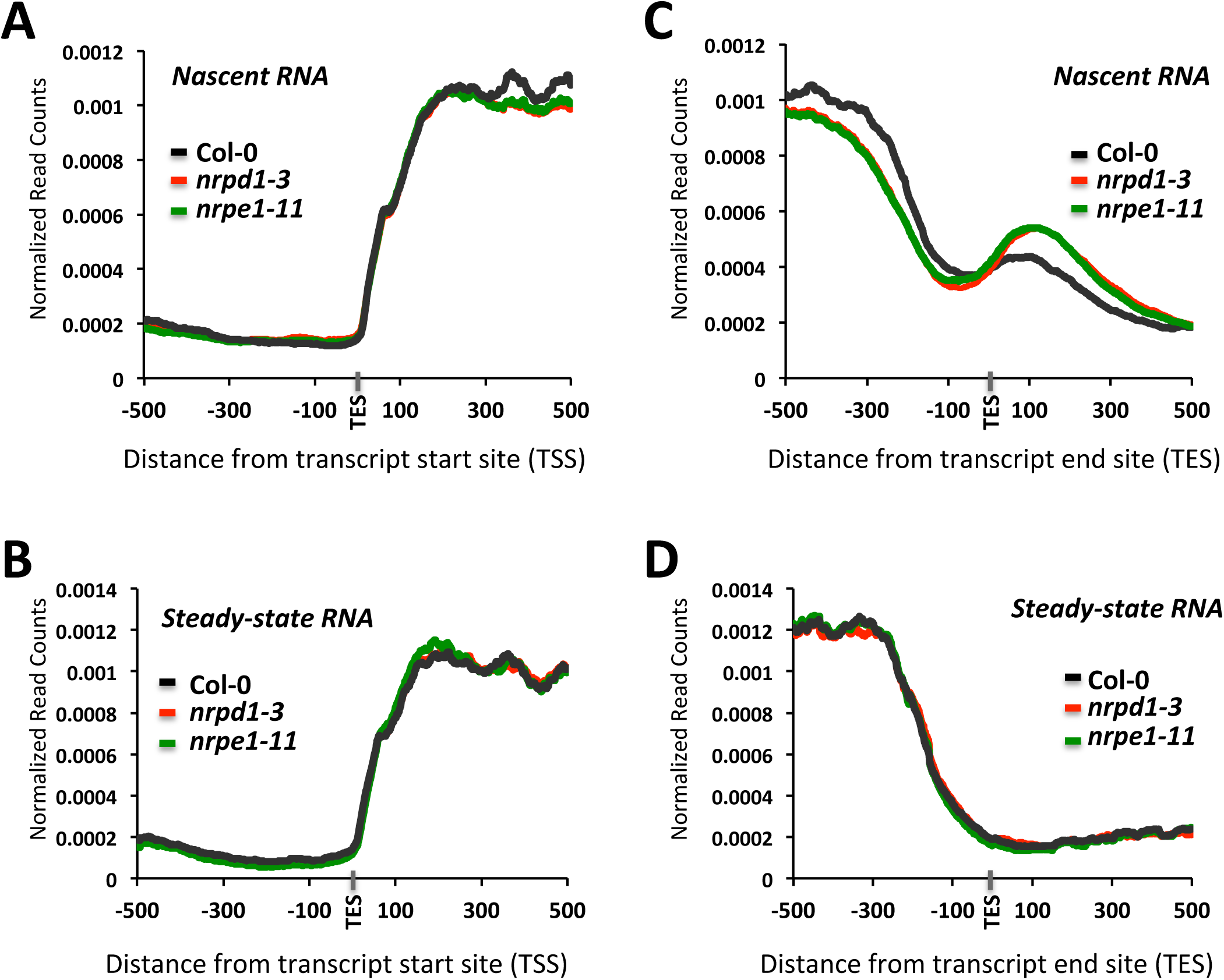
Loss of Pol IV or Pol V results in increased NRO-Seq read density downstream of protein-coding gene transcript end sites. **(A,B)** Average read densities are plotted relative to the transcription start sites (TSS) of 27,859 genes for nascent RNA **(A)** or steady-state RNA **(B)** isolated from Col-0, *pol IV* or *pol V* mutant plants. **(C)** Nascent RNA read-density plotted relative to transcript end sites (TES). **(D)** As in **(C)** but for steady-state RNA.

Loss of Pol IV or V activity does not affect RNA Pol II read density profiles near TSSs for either nascent RNA (Figure 1A, compare black with red and green) or steady-state RNA (Figure 1B), suggesting that RNA polymerases IV and V exert no detectable influence on Pol II transcription initiation at protein-coding genes on a genome-wide scale.

Analysis of the nascent RNA (NRO) read distribution in the 1 kb region surrounding the transcript end sites (TESs) revealed a gradual decrease in read abundance prior to the TES, followed by a small increase in reads ∼100 bp post-TES (Figure 1C, **in black**). This read density peak just downstream of TES in the nascent RNA population is not observed for steady-state RNA (Figure 1D, **in black**), suggesting that post-TES RNA is removed by cleavage at the polyA addition site and thus absent from mature mRNAs. Interestingly, a further increase in nascent transcript abundance occurs downstream of TESs in *pol IV* or *pol V* mutant plants, suggesting increased in Pol II occupancy, possibly due to delayed termination or Pol II release (Figure 1C).

To determine if specific genes have dramatically altered Pol II occupancy downstream of TESs in *pol IV* or *pol V* mutants, we calculated a transcript end site - associated termination index (TES-TI) for each gene as the number of total reads in the first 500 bp downstream of the TES (interval 2) divided by the number of total reads in the 500 bp preceding the TES (interval 1) (Figure 2A). Approximately 12% of the 18,419 protein coding genes analyzed exhibit a two fold or greater increase in their termination index in *pol IV* (*nrpd1-3*) or *pol V* (*nrpe1-11*) mutants, compared to wild-type (Figure 2B). Profiles for these genes, plotted as in Figure 1, are shown in Figure 2C. A strong concordance (∼80% overlap) is observed for genes affected in *pol IV* or *pol V* mutant plants (Figure 2B, **Venn diagram**). A set of 1312 genes, affected at least two fold by both *pol IV* and *pol V*, was chosen for all subsequent analyses. Loss of Pol IV or Pol V activities did not cause detectable lengthening of mature mRNAs for these genes, suggesting that RNA processing still occurs (Figure 2D), as illustrated in the gene browser profiles shown in Figure 2E.

**Figure 2:**
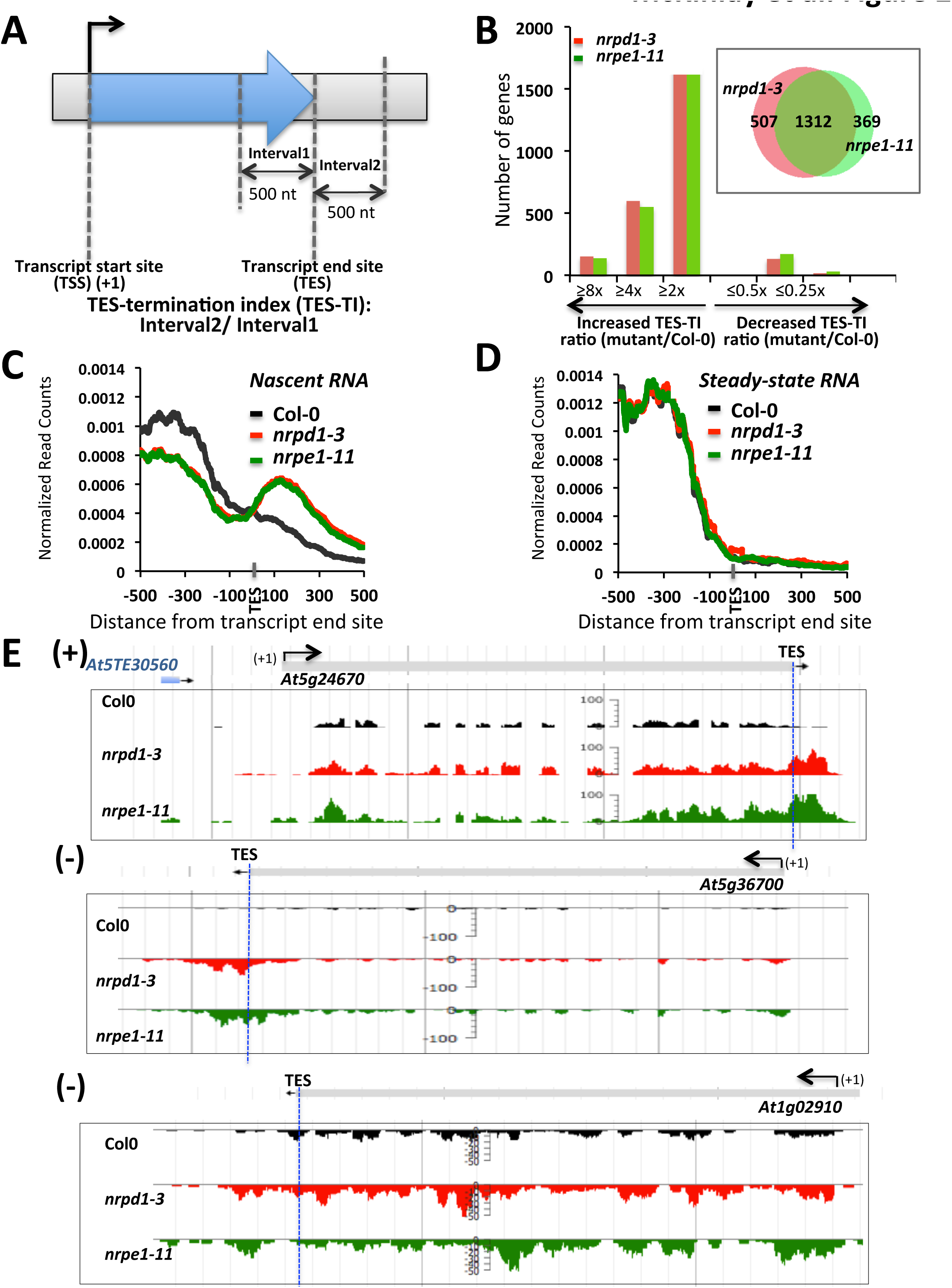
Genes with increased read density downstream of their transcript end sites (TES) in *pol IV* or *pol V* mutant plants. **(A)** Schematic for calculating the TES-termination index (TES-TI). **(B)** Histogram showing the number of genes with increased or decreased TES-TI values in *pol IV* or *pol V* mutant plants, relative to wild-type (Col-0) plants. **(Insert in B)** Venn diagram showing the overlap between genes with 2 fold or more increase in post-TES read density in *pol IV* (*nrpd1-3,* in red) or *pol V* (*nrpe1-11,* in green) plants relative to Col-0 plants. **(C)** Nascent RNA read-density plotted relative to TESs for the 1312 genes affected by both Pol IV and Pol V (see Venn diagram of panel **B**). **(D)** As in **(C)** but for steady-state RNA. **(E)** Examples of protein-coding genes with increased read density downstream of the TES (marked with blue dashed vertical line) in *pol IV* (*nrpd1-3*) or *pol V* (*nrpe1-11*) mutants, displayed using JBrowse.

To further examine whether changes in *pol IV* or *pol V* mutants affect qualitative or temporal aspects of 3’ end processing, we performed circular RT-PCR analyses at two loci, *At5g24670* and *At5g36700*, using primers that allow discrimination between full-length processed transcripts and transcripts that extend post-TES (Figure 3A). Longer circularized transcripts were detected in *pol IV* and *pol V* mutant plants, compared to wild-type Col-0 plants (Figure 3B, Suppl. Figure 3). Cloning and sequencing of these PCR products revealed that the majority of clones support the annotated gene models, but for the *At5g24670* transcript 14% of the cloned cDNAs in *nrpd1* and 24% of the cDNA clones in *nrpe1* include sequences downstream of the annotated TES (Figure 3C). Combined with our observation that steady-state mRNAs do not display changes in their 3’ ends, this suggests that 3’ end processing may be less efficient in *pol IV* or *pol V* mutants.

**Figure 3:**
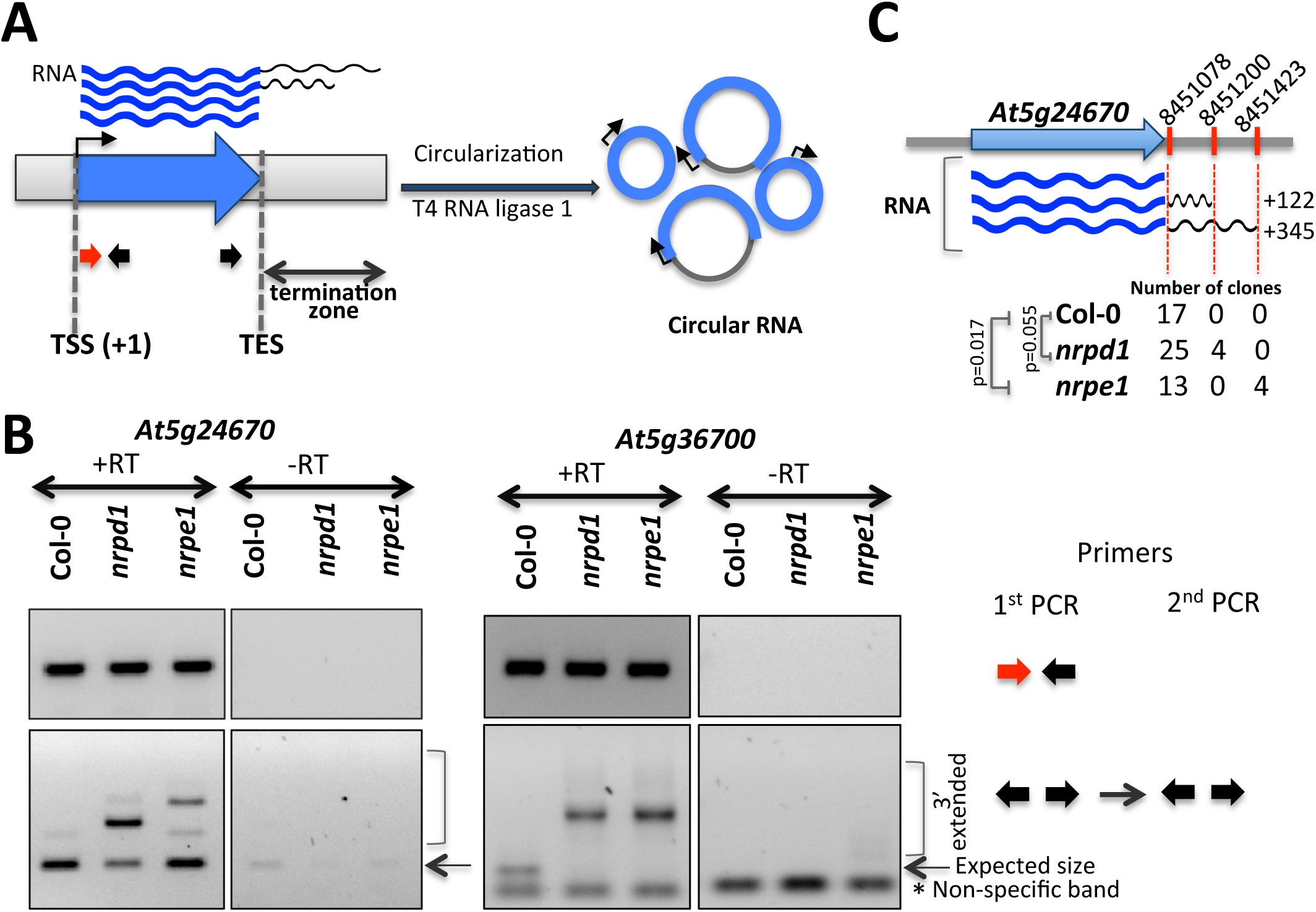
Transcripts with increased read density post-TES in *pol IV* or *pol V* mutant plants exhibit a longer termination zone. **(A)** Schematics of the circular RT-PCR approach used for independent confirmation of transcript ends. **(B)** Circular RT-PCR results for *At5g24670* and *At5g36700*. Top panels show PCR products obtained using primers placed near transcription start sites (TSS). Bottom panels show both mature and 3’-extended PCR products. Primers for each PCR reactions are color-coded as in **(A)** and are shown on the right. **(C)** Sequencing results for cloned *At5g24670* PCR products (from **B**, bottom panel). Number of clones supporting transcript ends at the marked (in red) genomic positions are shown for Col-0, *nrpd1,* and *nrpe1* plants. Numbers on the right represent observed increase (in nt) in the length of the transcripts relatively to mature transcripts. Significance has been tested using one-tailed Chi-square test.

### Pol II occupancy correlates with nuclear run-on signal intensity

Using chromatin immunoprecipitation (ChIP) of exonuclease-trimmed chromatin-protein complexes (ChIP-exo), we examined Pol II association with genes displaying increased (>2x) post-TES nascent RNA-seq read density in *pol IV* or *pol V* mutants (Figure 4). The ChIP-exo method allows high resolution mapping of DNA binding factors due to the exonuclease trimming of sequences not protected by the bound protein^43^. Consistent with NRO-seq results, the ChIP-exo sequence data revealed increased Pol II association with gene 3’ ends in *pol IV* mutants (Figure 4A), compared to Col-0 wild-type, with a strong peak downstream of the TESs. In contrast, no differences in RNA Pol II occupancy was observed near the transcription start sites for the same set of genes (Figure 4B), suggesting an asymmetric affect at gene 3’ ends.

**Figure 4:**
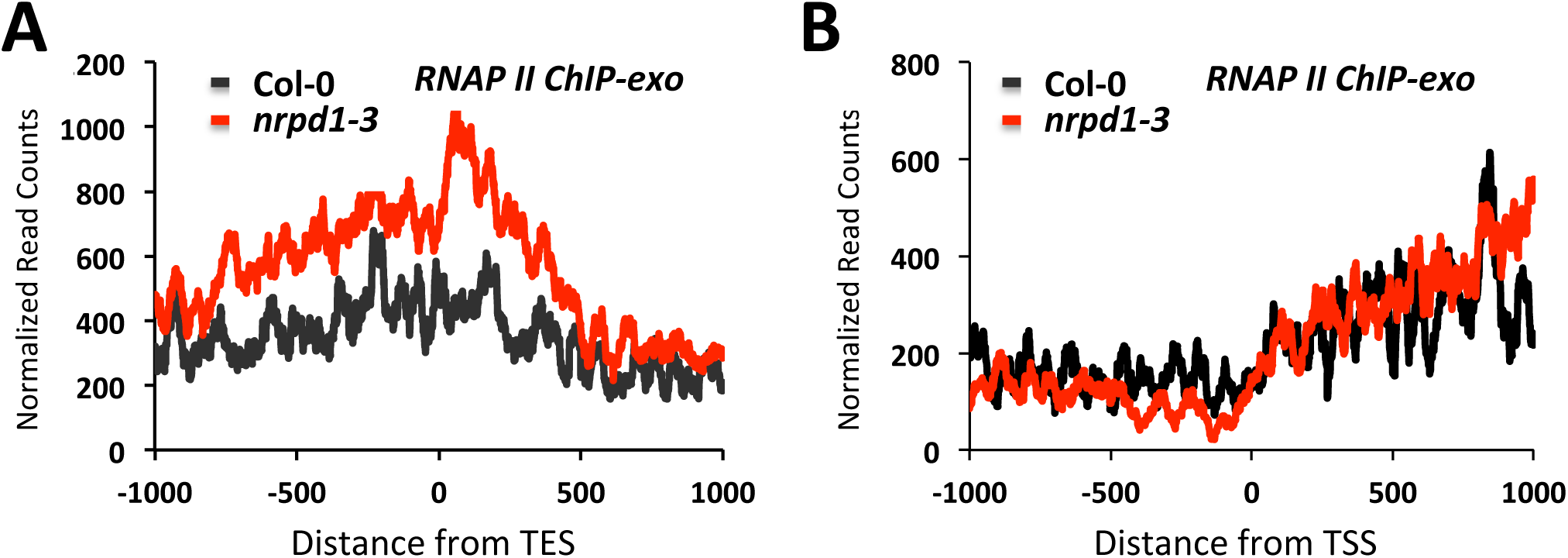
Pol II accumulates post-TES in *pol IV* mutant plants. **(A)** RNA Pol II ChIP-exo reads are plotted relative to transcript end sites (TES) for genes with increased post-TES read density in *pol IV* or *pol V* mutants. **(B)** As in **(A)** but reads are plotted relative to TSSs.

### Genes with Pol IV and V – affected transcription termination associate with RdDM pathway components

We examined published Pol IV^44^, Pol V^17^, and AGO4^45^ ChIP-seq datasets to see if these proteins are enriched near the 3’ ends of genes that show increased Pol II occupancy post TES in *pol IV* or *V* mutant plants. Interestingly, for 60% of the 1312 genes with increased post-TES read density, Pol IV, Pol V, or AGO4 binding sites overlap the 1 kb region surrounding their TESs, with Pol V being present at 50% of these genes (Figure 5A). Analysis of Pol V ChIP-seq^17^ read distribution in wild-type (Col-0) plants revealed Pol V association primarily upstream of TESs, with an additional peak, compared to background (*nrpe1-11*), 150-250 bp downstream of the TES (Figure 5B). Analysis of Pol IV ChIP-seq^44^ data did not reveal evidence for a similar pattern of enrichment (Suppl. Figure 4).

**Figure 5:**
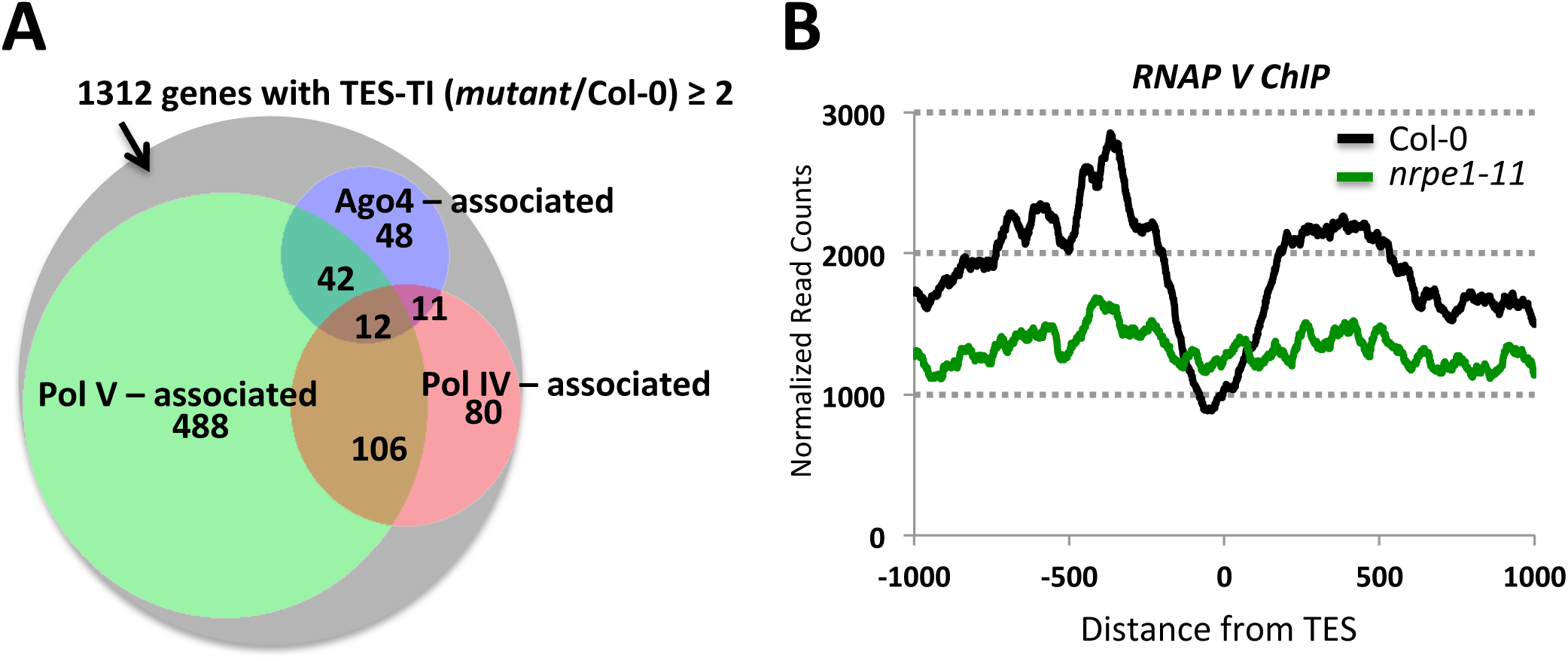
Correspondence between genes with increased read density post-TES and RdDM pathway protein chromatin interaction. **(A)** Venn diagram showing the number of genes with increased read density post-TES that exhibit enrichment (≥ 2 fold) for Pol IV, Pol V, or AGO4 within 2 kb regions centered at their TESs. **(B)** Pol V ChIP-seq read density plotted for genes with increased nascent RNA reads post-TES in *pol IV* or *pol V* mutant plants.

Fewer than 1% (12 out of 1312) of the analyzed genes displayed overlapping Pol IV, Pol V, and AGO4 occupancy post-TES, suggesting that loci with increased post-TES NRO signal are not likely subjects of RNA-directed DNA methylation. To further confirm this statement we examine whether 23-24 nt siRNAs map near TESs of genes affected by the loss of Pol IV or Pol V. Only 4.2% of affected genes overlap with 23-24 nt and 3.5% with 21-22 nt siRNA reads within a 1 kb region surrounding the TESs (Suppl. Figure 5A). Similarly, no correlation between post-TES read density and CHH methylation was detected in *rdd* (*ros1-3 dml2-1 dml3-1*) triple mutant^46^ that lacks nearly all DNA demethylation activity (Suppl. Figure 5B). This suggests that effect of Pol IV and Pol V on Pol II transcription termination is unlikely to be due to RNA-directed DNA methylation. Interestingly, CG methylation profile mimics Pol V occupancy at genes that show increased Pol II signal post TES in *pol IV* or *V* mutant plants (Suppl. Figure 6). Consistent with previous report^47^, CG methylation might guide Pol V to Pol II transcript end sites. Whereas lack of CHH methylation and siRNA accumulation at these sites suggests that Pol V is not transcriptionally engaged.

### Pol II accumulation in the post-TES region is increased for convergent gene pairs

Interestingly, convergent gene pairs were significantly enriched among genes with increased post-TES read density in *pol IV* or *pol V* mutant plants (Figure 6A). Furthermore, these genes tend to have neighboring genes that initiate or end within 100 bps downstream or 100 bps upstream of their TESs (Figure 6B).

Regulation of transcription termination of convergent gene pairs might be an important mechanism to modulate levels of natural anti-sense transcripts (NATs) in cells as the base-pairing between sense and antisense transcripts can form dsRNAs, which can be then processed into *cis*-NAT-derived siRNAs (nat-siRNAs) via unique Pol IV-dependent pathway and lead to down-regulation of transcript levels though mRNA cleavage^48,49^. Analysis of antisense and sense reads revealed that both were increased within 1 kb region surrounding TESs for genes with increased Pol II occupancy post-TES in *pol IV* or *V* mutant plants (Suppl. Figure 7A). However, we didn’t detect the expected increase in numbers of neither 21-22 nt nor 23-24 nt sRNA clusters (Suppl. Figure 7B) nor did we detect statistically significant changes in transcript abundance for either gene in the convergent gene pairs (Suppl. Figure 7C), although it is possible that a specific abiotic or biotic stress condition might be required. Indeed, gene ontology analysis revealed that genes with increased post-TES read density are enriched for transcripts involved in response to abiotic stress (*P* = 8.0 × 10^−6^) (Suppl. Figure 8).

**Figure 6:**
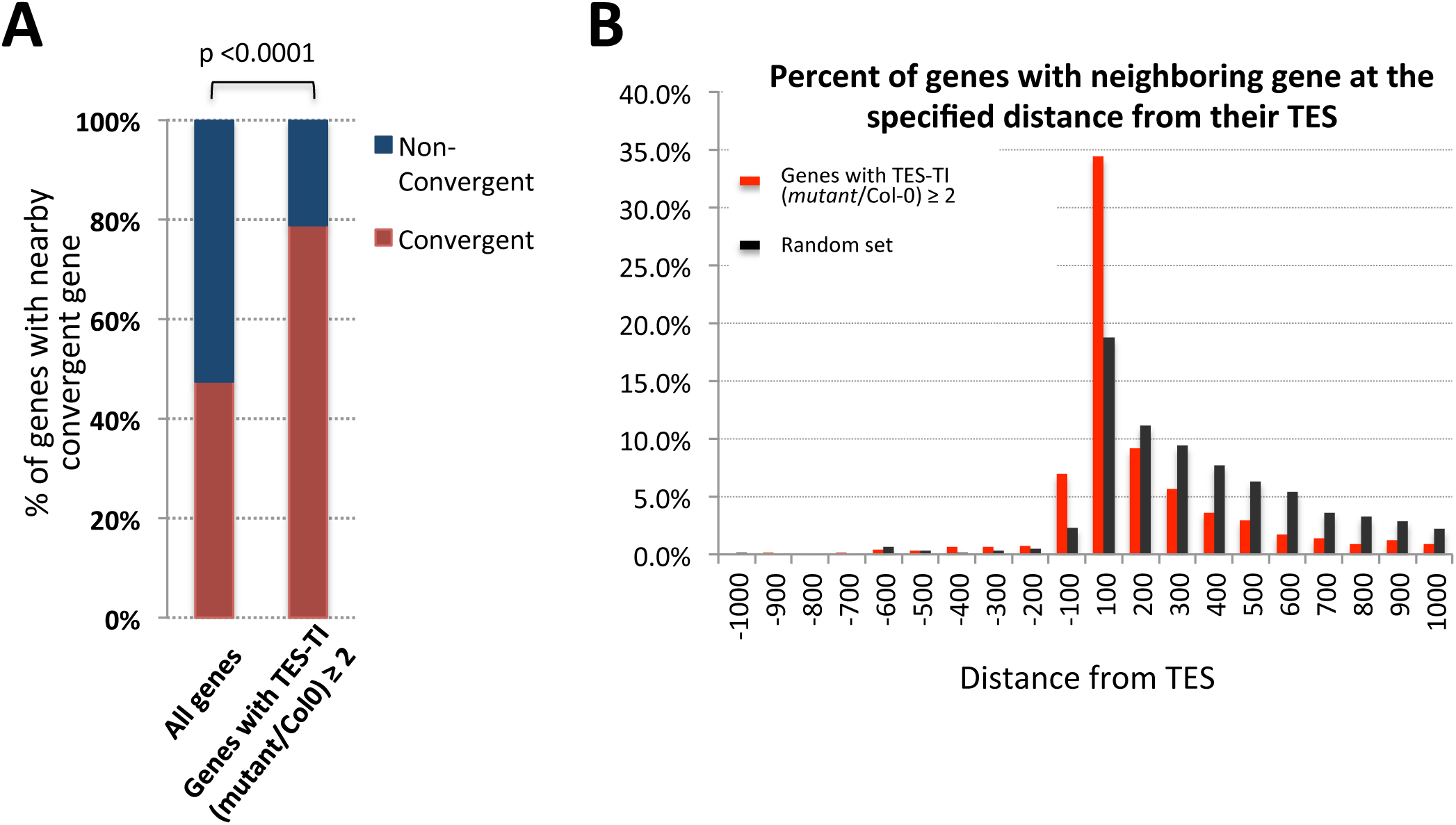
post-TES read accumulation is more prevalent among convergent gene pairs. **(A)** Histogram showing the percent of convergent genes among all protein-coding genes in *Arabidopsis thaliana* versus genes displaying increased post-TES read density in *pol IV* or *pol V* mutant plants. *P*-value was calculated using Fisher’s exact test. **(B)** Histogram showing the percent of genes with increased post-TES read density in *pol IV* or *pol V* mutant plants that have the nearest neighbor gene initiating or ending at various distances (x-axes) from their transcript end sites (TES).

## Discussion

Nuclear run-on combined with high-throughput sequencing (NRO-Seq)^36^ and RNA-Seq^50^ revealed that Pol IV and Pol V affect Pol II occupancy downstream of polyA addition sites (TESs), for approximately 12% of protein-coding genes. Consistently, Pol IV or Pol V interaction sites include the TES regions for genes that display increased NRO signals downstream of TESs in *pol IV* or *pol V* mutant plants. However, for the majority of these genes there is no enrichment for 24-nt siRNAs and/or changes in CHH methylation, a hallmark of Pol IV and Pol V - dependent RdDM. Thus, any influence of Pol IV and Pol V on Pol II transcription termination is likely independent of DNA methylation.

Pol II elongation rates can affect co-transcriptional processes such as transcription termination resulting in many RNA polymerase molecules continue transcribing post-TES^51^. Interestingly, genes with high post-TES NRO signals in *pol IV* or *pol V* mutant plants exhibit not only higher Pol II occupancy near TES (Figure 4), but also significantly higher rates of transcription, compared to all protein-coding genes (Suppl. Figure 9 and Figure 2E). Elongation rate could be influenced by various factors that can modulate Pol II transcription efficiency directly (e.g. elongation factors) or indirectly, by changing chromatin organization (chromatin remodelers). Pol V transcription can recruit the SWI/SNF chromatin remodeling complex, such that a loss of Pol V transcripts correlates with changes in nucleosome positioning^52^. Whether or not Pol IV can similarly affect chromatin states has not yet been determined. However, CLASSY1, a putative ATP-dependent DNA translocase, has been implicated in Pol IV function and speculated to affect nucleosome repositioning, similar to SWI/SNF^53,54^. Thus, Pol IV or Pol V – mediated chromatin modifications near Pol II TESs may also influence Pol II termination or release.

Interestingly, convergent gene pairs tend to show the highest post-TES NRO signals in *pol IV* or *pol V* mutant plants. Convergent transcription can result in head-to-head collisions of transcribing Pol II enzymes^55^. Though such events are expected to be rare, Pol II collision doesn’t result in polymerase dissociation from the template but leads to transcriptional arrest and gene blockage^55^. To date, not much is known about how cells deal with collision-induced transcription blocks. However, Pol II ubiquitination-mediated degradation has been shown to be involved in clearing Pol II from the region between convergent genes, such that lack of the yeast RNA Pol II ubiquitination/degradation protein ELC1 results in a peak of RNA Pol II density between convergent genes^55^. A possibility is that Pol IV or Pol V occupancy at intergenic DNA may reduce RNA polymerase II collisions by slowing or dissociating Pol II, thus allowing more efficient cleavage at polyA sites and subsequent termination (Figure 7). Consistently, we observed lower Pol II occupancy just upstream of TES (Figure 1C and Figure 2C) as well as higher levels of 3’ unprocessed transcripts (Figure 3) in *pol IV* or *pol V* mutant plants.

**Figure 7:**
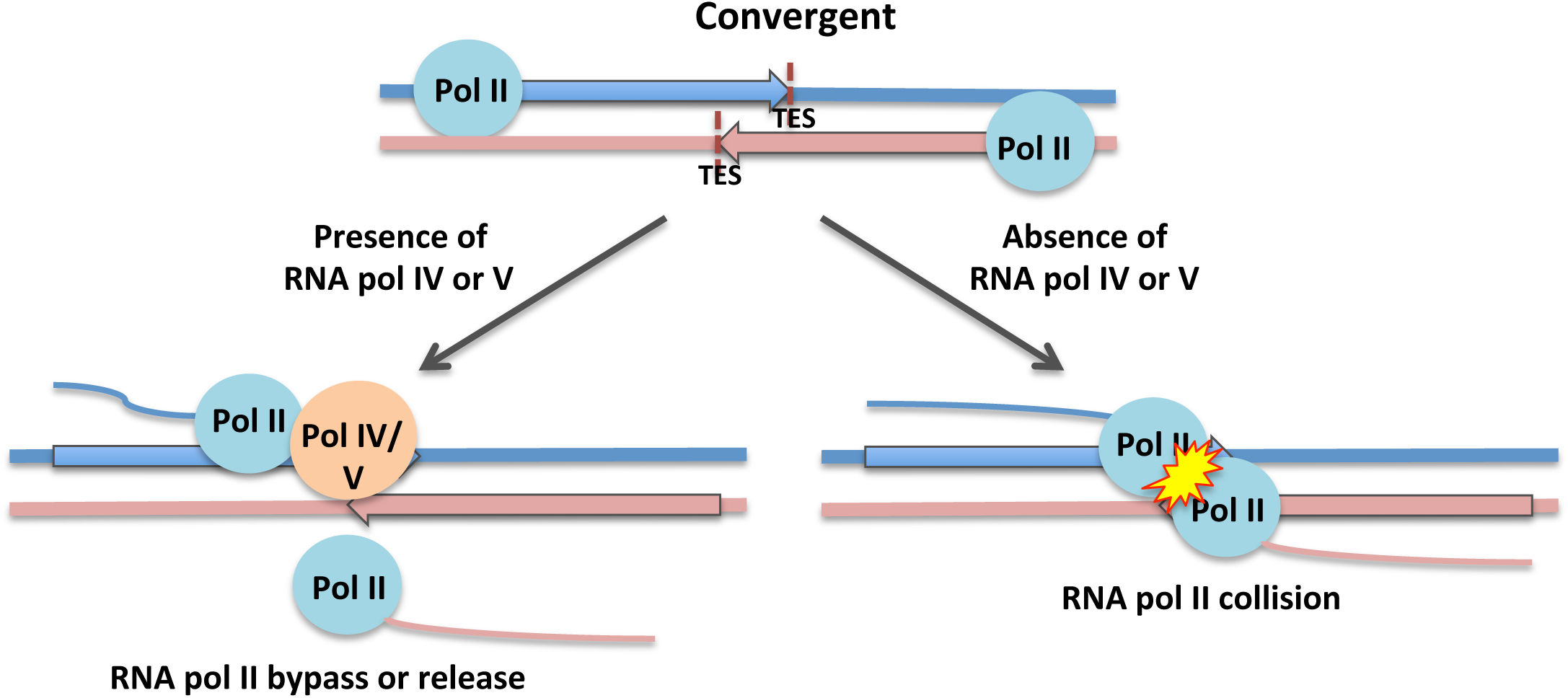
Model for RNA Pol IV or Pol V involvement in RNA Pol II transcription termination. Pol IV or Pol V may impede Pol II to promote transcript cleavage and release at Pol II convergent genes. In the absence of Pol IV or Pol V, Pol II undergoes head-to-head collision and pausing.

Pausing or slowing down of Pol II near the TESs during transcription termination is thought to facilitate co-transcriptional splicing^51,56,57^. Interestingly, splicing was much less abundant in *pol IV* or *pol V* mutant plants, compared to wild-type (Suppl. Figure 10), suggesting that Pol IV or Pol V affect pre-mRNA processing at the level of transcription. Alternatively, or in addition, Pol IV or Pol V might facilitate recruitment of splicing factors^58^. For example, a U4/U6 snRNP-associated protein, RDM16, is enriched at Pol V target loci, suggesting that RDM16 might be recruited by Pol V and/or by the nascent Pol V transcripts^59^.

## Materials and Methods

### Plant material

*A. thaliana* Col-0 and *nrpd1-3* and *nrpe1-11* have been described previously^2,15^. Plants were cultivated at 22°C under long-day conditions (16 h day/8 h night).

### Nuclei isolation

Nuclei were isolated from 3 week old Col-0, *nrpd1-3* and *nrpe1-11* plant seedlings as described previously^60^ with a few modifications. Briefly, 3 week-old plants were harvested and washed in ice-cold water. Then plants were submerged in cold diethyl ether for 5 min and washed twice in ice-cold water. Homogenization of plant material was performed in 100 mL of nuclei isolation buffer (1M sucrose, 50 mM Tris-HCl [pH 7.2], 5 mM MgCl, 5 mM KCl, 10 mM 2-mercaptoethnol, 0.2 mM PMSF) with a motorized homogenizer (Fisher Scientific Powergen 700). Homogenate was filtered through 1 layer of Miracloth and 1 layer of 50 um of nylon mesh. After centrifugation at 14,000 g for 15 min at 4°C the pellet was resuspended in 10 min of nuclei isolation buffer and prepared for loading into discontinuous Percoll gradient (15%, 30%, 45%, and 60%) by adding one volume of “100% Percoll solution” (34.23 g sucrose, 5 mL 1 M Tris-HCl [pH 7.2], 0.5 mL 1 M MgCl_2_, 0.5 mL 1 M KCl, 34 uL 2-mercaptoethnol, 100 uL 2 mM PMSF and Percoll to 100 mL) to 19 volumes of nuclei. Gradients were centrifuged at 500 g for 10 min, then 7,500 rpm for 20 min. Nuclei were harvested from 30%/45% and 45%/60% boundaries and pooled. Nuclei were diluted with 5 volumes of nuclei isolation buffer, mixed by inversion and collected by centrifugation at 1,500 g, 10 min, 4°C. Second wash of nuclei was performed with 25 volumes of nuclei isolation buffer. Final resuspension of nuclei was in 1 mL of nuclei storage buffer (50 mM HEPES [pH 7.2], 5 mM MgCl_2_, 5 mM KCl, 2 mM DTT, 0.2 mM PMSF, 50% glycerol). Aliquots were frozen in liquid nitrogen and stored at −80°C.

### Nuclear run-on (NRO)

NRO reactions were performed as described previously^36^ with a few modifications. Briefly, 10 *in vitro* transcription reactions (each containing190 uL of nuclei, 60 uL 5x Txn buffer (100 mM Tris-HCl, [pH 7.7], 6 uL 0.1 M DTT, 5 uL RiboLock, 12 uL Biotin-16-UTP (10 mM), 16 uL AGC (10 mM each)) were set up in parallel for each plant line examined and incubated at 30°C for 60 min. Total RNA was extracted using Trizol method, treated with Turbo DNase I. An aliquot of total RNA was used for total RNA library contruction (see Materials and Methods). Biotinylated RNA was precipitated from total RNA with Dynabeads MyOne Streptavidin C1 (Invitrogen, Calsbad, CA). Binding was performed in 5mM Tris-HCl [pH 7.7], 0.5 mM EDTA and 1 M NaCl for 20 min at 42°C and 2 hours at room temperature with rotation. After washes, RNA was eluted from beads by adding 50 uL of 10 uM EDTA/95% formamide and incubation at 65°C for 5 min.

### RNA library construction

Libraries from total and eluted biotinylated RNA were prepared using the Illumina TruSeq Stranded mRNA Sample Prep Kit. No mRNA or rRNA depletion steps were performed. Libraries were subjected to 50 bp paired-end sequencing.

### exoChIP library construction

Chromatin cross-linking, isolation and shearing were carried out as previously described with slight modifications^15^. Briefly, three grams of three-week old leaf tissue was suspended in 35 ml SH Buffer (100 mM sucrose, 20mM HEPES-NaOH) containing 1% formaldehyde and placed under vacuum for 55 minutes. Cross-linking was stopped by the addition of 1.25 ml 2M Glycine. Leaf tissue was washed three times with water. Tissue was ground in mortar with pestle and suspended in 15 ml Honda buffer (0.44M Sucrose, 1.25% Ficoll, 2.5% Dextran T40, 20mM HEPES-NaOH pH7.4, 10mM MgCl_2_, 0.5% Triton X-100, 5mM DTT, 1mM PMSF, 1% Plant Protease Inhibitor Cocktail). Slurry was passed through Miracloth and centrifuged at 2,000 x g for 15 minutes at 4C. The pellet was resuspended in 1 ml Honda buffer and centrifuged as above. Again, the pellet was resuspended in Honda buffer, centrifuged as above and the pellet was resuspended in 575 microliters of Nuclei Lysis Buffer (50mM Tris-HCl pH8, 10mM EDTA, 1% SDS, 1mM PMSF, 1% Plant Protease Inhibitor Cocktail). Chromatin was sheared using a Bioruptor sonicator (Diagenode) to generate fragment sizes between 200 and 600 bp. Sheared chromatin was centrifuged for 10 minutes at 4C at 12,000 x g. The supernatant was removed and diluted ten fold with ChIP Dilution Buffer (1.1% Triton X-100, 1.2mM EDTA, 16.7mM NaCl, 16.7mM Tris-HCl pH8). Aliquots containing approximately 6 mg of chromatin were shipped to Peconic, LLC for ChIP-exo sequencing. ChIP-exo was carried out as described in^61^. Five microliters of the 4H8 monoclonal antibody (Cell Signaling Technology), which recognizes the Pol II CTD with a preference towards the phosphoS5 form, was used for immunoprecipitation.

### Circular RT-PCR

Total RNA was extracted using Trizol reagent. 10 ug of RNA was decapped by treatment with Tobacco Acid Pyrophosphatase (TAP) (Epicentre) to provide 5’-monophosphorylated terminus that was ligated to a 3’-hydrohylated terminus by T4 RNA ligase 1 (NEB, 10,000 U/uL) according to manufacture instructions. After incubation at 37°C for 1 hour, enzymes were heat inactivated at 65°C for 15 min and reactions were cleaned up and concentrated with Zymogen RNA Clean and Concentrator kit. The resulted RNA was treated with Turbo DNase I, reverse-transcribed with gene-specific reverse primer. Two rounds of PCR amplification was followed by PCR clean up (Qiagen PCR clean up kit) and DNA cloning into pGEMT easy vector according to manufacturer’s instructions. Individual colonies were grown and used for DNA extraction and sequencing.

Primers

AT5G36700_F1 ccaaaagaagcggaaacaga

AT5G36700_R ccgccacagaaagaagaaga

AT5G36700_F2 atttgcagtgcctttgtgtg

AT5G24670_F1 tcatcattgttgtttccttctca

AT5G24670_R atgagcggagcagctactgt

AT5G24670_F2 gcgcttgtgcatcaaagaat

### Sequence processing

Reads were adapter trimmed and quality filtered (q20) using trimmomatic and then mapped to TAIR10 genome reference using tophat. All transposable elements, tRNAs, and ribosomal RNAs, as well as genes in the mitochondria and chloroplast genomes were excluded from analysis. The reads were further filtered to remove low complexity reads (with the entropy of 2 or less) using custom Perl scripts. Only read alignments mapped concordantly (e.g. with correct orientation and separated by less than 7,000 bps from each other) were selected for subsequent analysis. The 7,000 bp distance cut off was based on the 99.9% intron length in the TAIR10 annotation. Read data was converted into coverage information in the form of strand specific bed files. Cumulative coverage data was determined in a strand-specific way from the 5’ and 3’ ends of genes as defined in the TAIR10 gene file. The read counts between samples were normalized based on the total number of reads mapped for each sample. *AT2G16586* was excluded from analysis due to it having an extremely high and variable coverage between samples.

### Read-through analysis

TES-termination index (TES-TI) was determined in the NRO and total RNA libraries by calculating ratios of read density 500 bp downstream of the TES to read density in the body of the gene 500 bp upstream of TES. For each gene, TES-TI was calculated in Col-0, *nrpd1-3* and *nrpe1-11* backgrounds.

### ChIP data processing

Among the annotated TAIR10 nuclear protein coding genes, pairs of consecutive genes with their 3’ ends facing each other and separated by at least 1kb were identified as convergent genes. Among these, any gene pairs overlapping with miRNA or tRNA or snRNA or snoRNA or rRNA or other non-coding RNA were excluded, resulting in 1,453 gene pairs (2,906 genes) that were used in the analysis. For each of these 2,906 genes, coverage information at each base position starting from 1000 nt upstream of the 3’ end into the gene body to 1000 nt downstream of the 3’ end was obtained and total coverage contributed by all 2,906 genes at base position was computed and plotted.

### Bisulfite sequencing analyses

Bisulfite sequencing reads were quality processed (q >/= 25) and adapter trimmed using Cutadapt version 1.9.1^62^. Cleaned reads were mapped to the Arabidopsis thaliana TAIR10 genome using Bismark version 0.16.1 default settings^63^. PCR duplicates were removed and methylation information for cytosines in the CHH context with a minimum 5 read coverage were used for further analyses. Differentially methylated regions (DMRs) that showed a minimum 10% increase in methylation levels in rdd relative to Col-0 were identified using the R package methylKit version 0.9.5^64^. Regions were defined using a 300 base pair sliding windows with a step size of 200 base pairs. The minimum coverage requirement for each window was 10 cytosines with at least 5 read coverage each and significance was defined as a q value less than 0.01. Overlapping DMRs were merged into a single region and genomic coordinates of rdd hyper-methylated regions (relative to Col-0) were used to assess overlap with regions flanking the 3’ end of genes of interest 500 basepairs upstream and 500 basepairs downstream. Accession numbers for Col-0 and rdd bisulfite sequencing data are GSM276809 and GSM276812, respectively^65^.

### sRNA sequencing analyses

Raw reads were adapter and quality trimmed and size selected using Cutadapt version 1.9.1. Reads were further filtered of all structural RNAs (tRNAs, rRNAs, snRNAs, and snoRNAs) and mapped to the *Arabidopsis* TAIR10 genome using ShortStack version 3.4 default settings^66^. sRNA counts for regions of interest were extracted from bam files using the ShortStack --locifile file function and 21-24nt siRNA clusters were defined using a minimum read coverage of 25 reads. The accession number for Col-0, *nrpd1-3*, and *nrpe1-11* sRNA sequencing data are SRR2075819, SRR2505369, and GSM2451982, respectively^6^.

### Data availability

Raw Illumina reads will be deposited to the NCBI Gene Expression Omnibus (GEO) upon acceptance for publication and will be available through http://www.ncbi.nlm.nih.gov/geo/.

## Acknowledgements

We thank present and former Pikaard lab members for valuable discussions and constructive criticisms. We thank Craig Pikaard for help with editing manuscript. We thank Jered Wendte for assistance with circular RT-PCR technique.

## Author Contributions

A.M. designed and performed the research; R.C. performed ChIP-exo experiments; D.B.R., R.P., J.M.W., and A.M. analyzed the data; A.M. wrote the paper; A.M., D.B.R. revised manuscript and figures.

---

**Suppl Figure 1:**
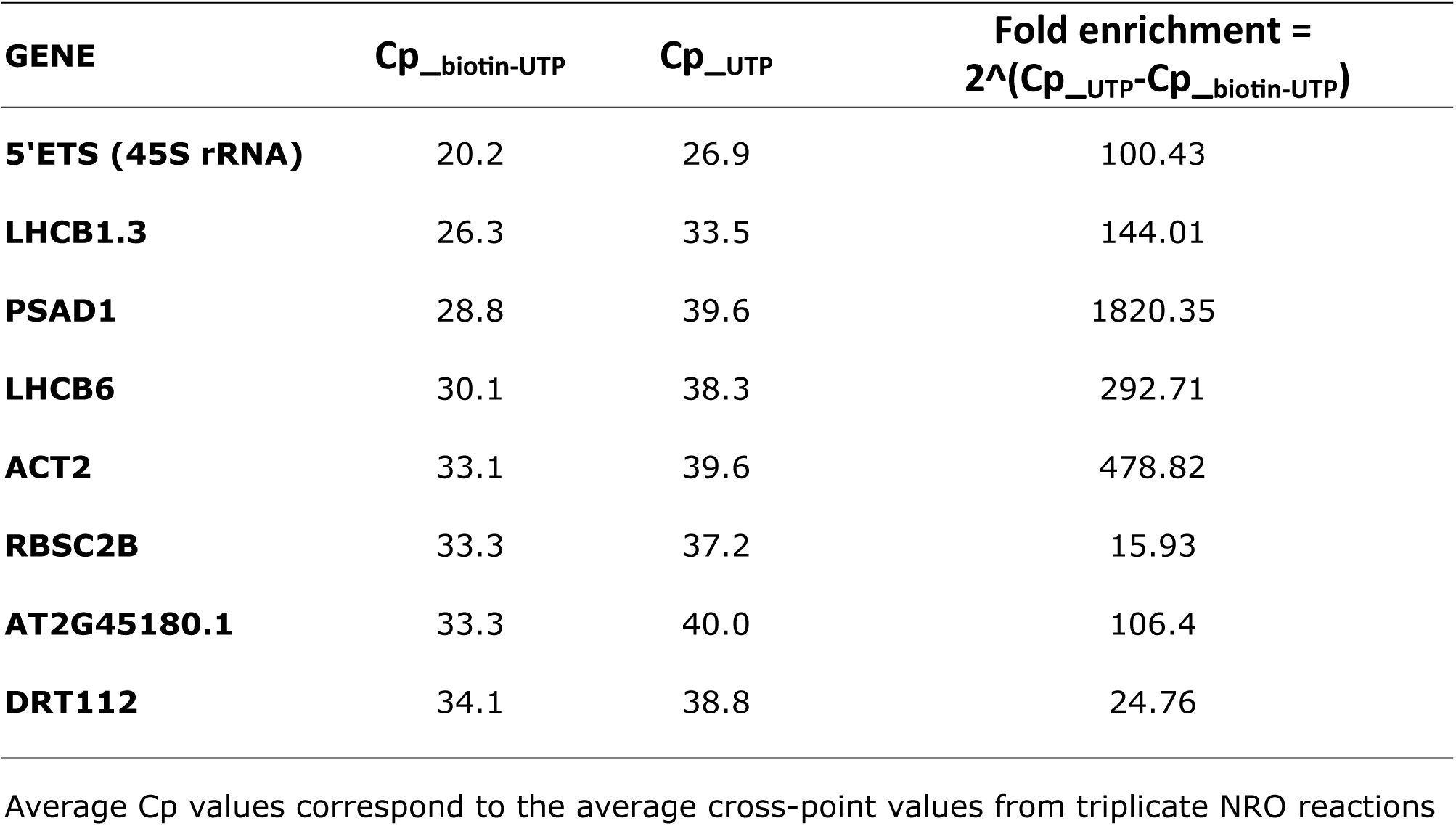
Specificity of nascent biotinylated RNA enrichment on streptavidin beads. Nuclear run-on (NRO) reactions were performed in triplicates with either biotin-UTP or with non-biotinylated UTP (as a negative control). Nascent RNA was captured on streptavidin beads, eluted and used in qPCR assays. The table shows averages of crossing point (Cp) values for several highly expressed *Arabidopsis thaliana* genes. Fold enrichment was calculated as the difference in Cp values between UTP samples and biotin-UTP samples: as Fold enrichment = 2^∧^(Cp__UTP_-Cp__biotin-UTP_).

**Suppl Figure 2:**
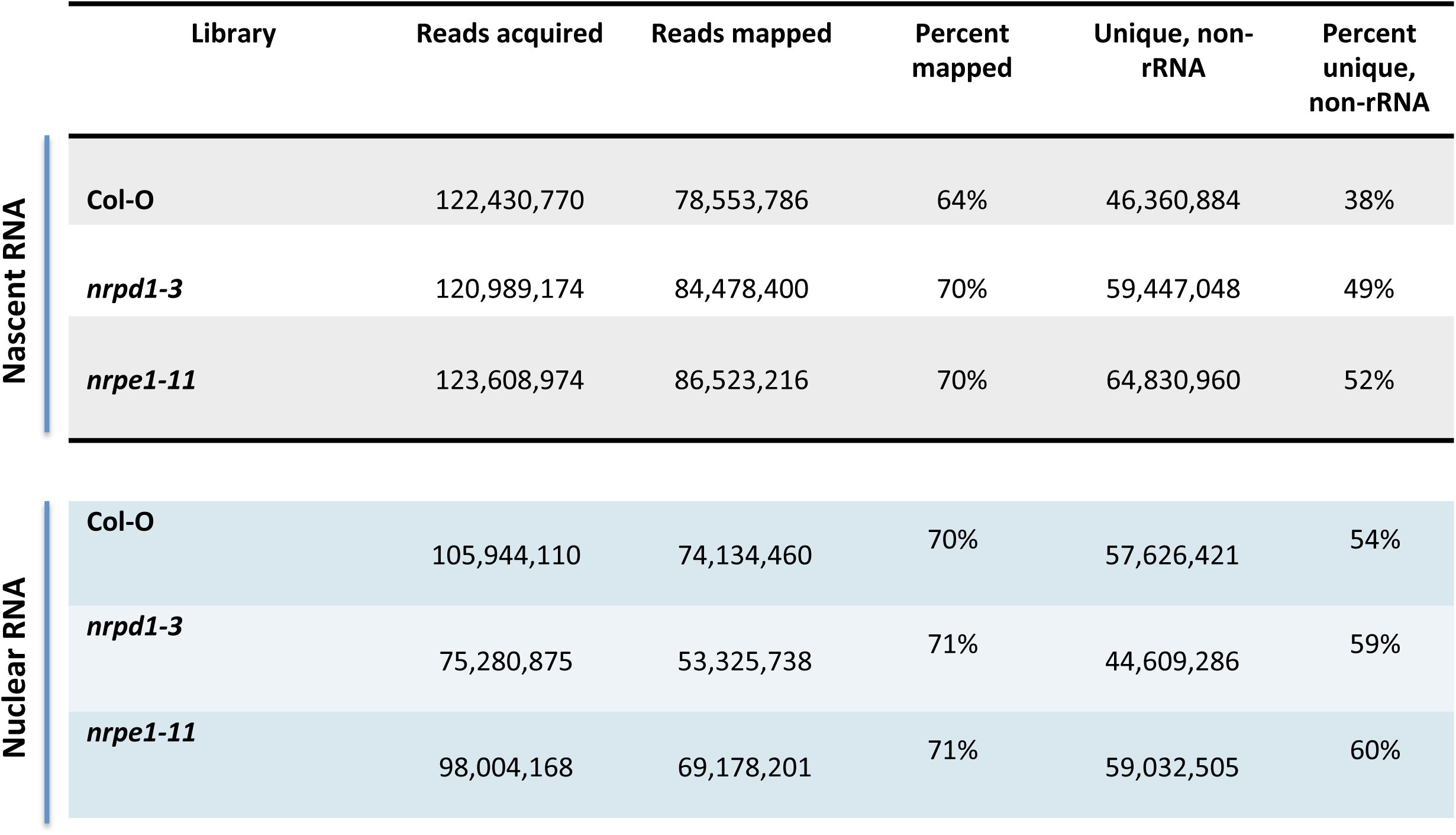
Sequencing data mapping statistics. Acquired sequence reads were mapped to TAIR10 RefSeq. All non-unique reads were filtered to obtain high-quality sequence reads for metagene analyses.

**Suppl Figure 3:**
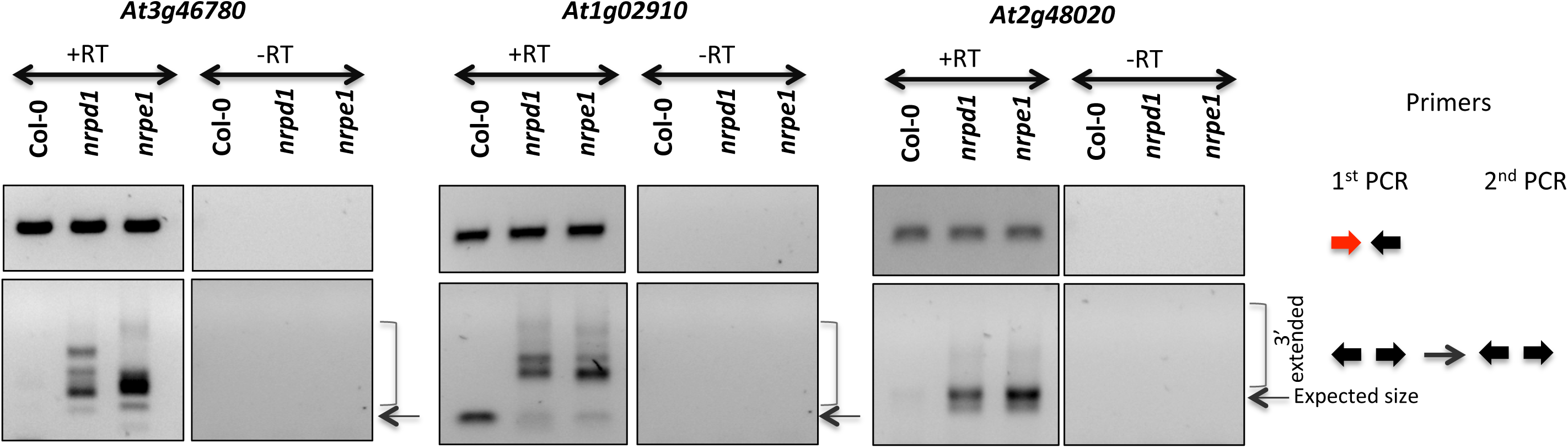
Circular RT-PCR results. Circular RT-PCR results for *At3g46780, At1g02910,* and *At2g48020*. Top panels show PCR products obtained using primers placed near transcription start sites (TSS). Bottom panels show both mature and 3’-extended PCR products. Primers for each PCR reactions are color-coded as in Figure 3A and are shown on the right.

**Suppl Figure 4:**
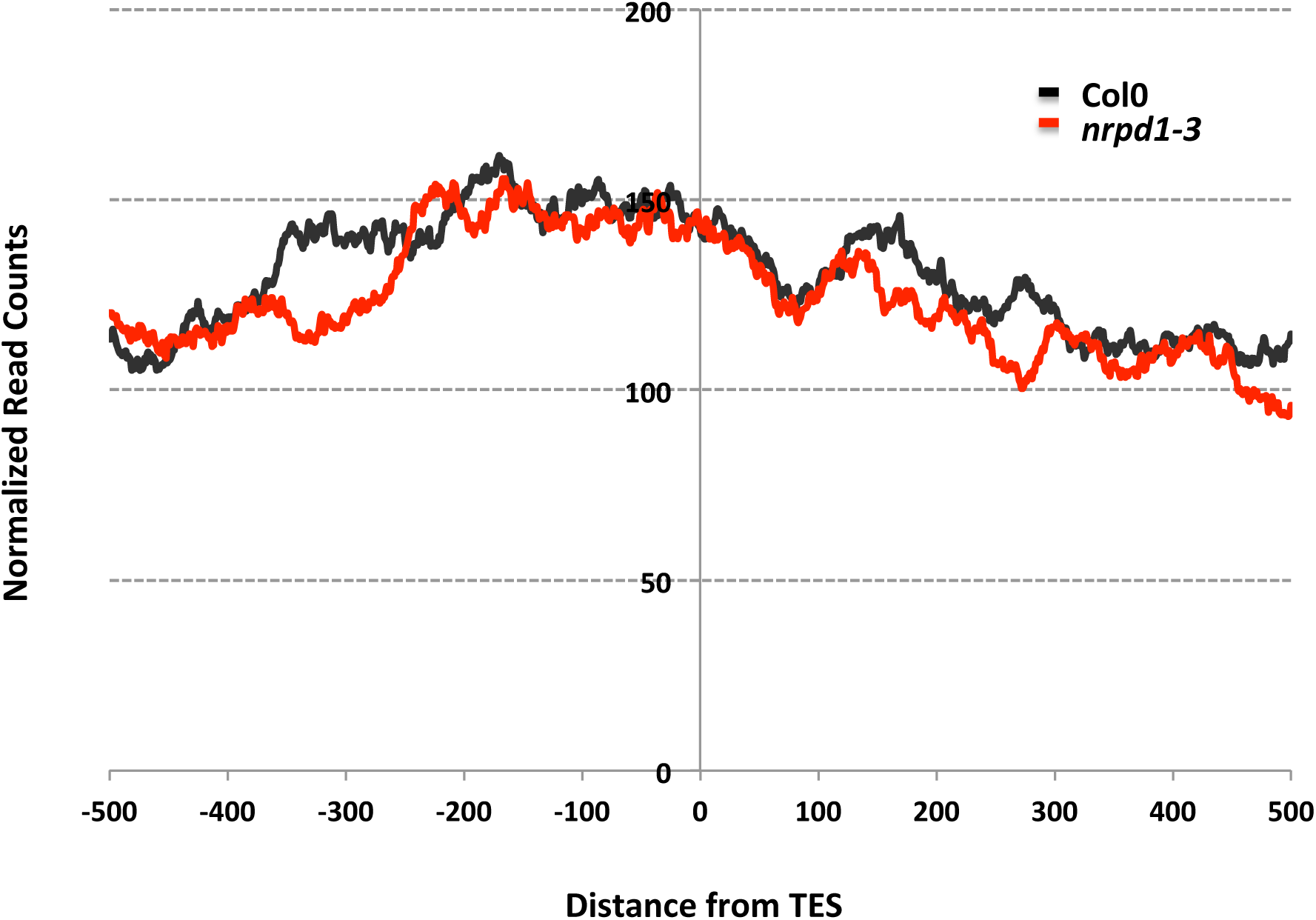
Pol IV ChIP-seq read density plotted for genes with increased nascent RNA reads post-TES in *pol IV* or *pol V* mutant plants.

**Suppl Figure 5:**
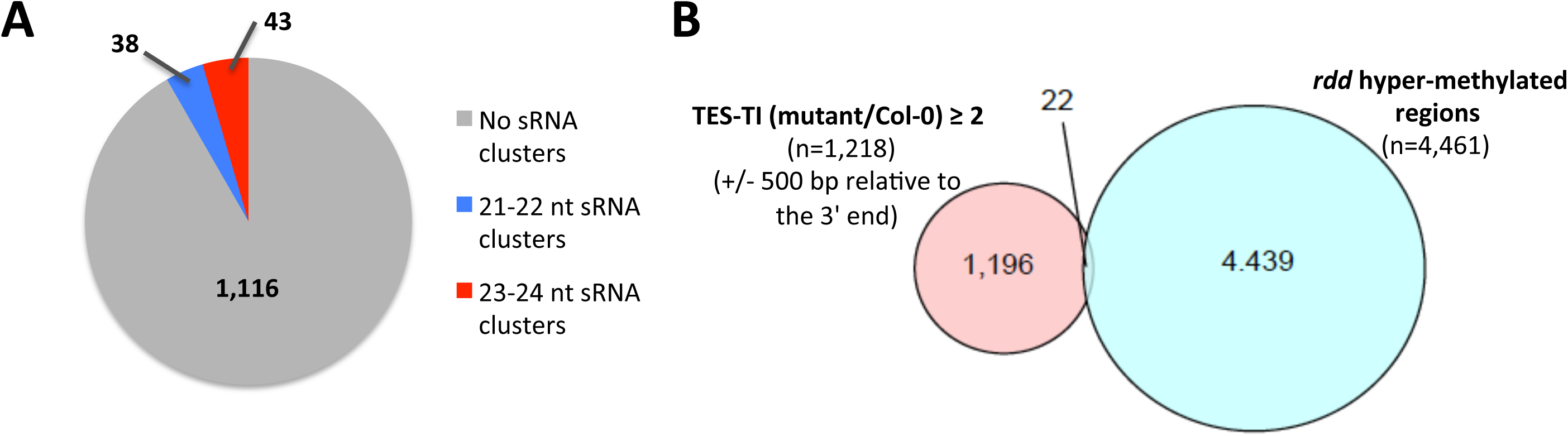
Increase in post-TES read density in *pol IV* or *pol V* mutants is independent of siRNA biogenesis and CHH DNA methylation. **(A)** The Pie chart shows number of genes with increased post-TES read density that overlap with 21-22 nt or 23-24 nt sRNA clusters within a 1 kb region surrounding the TESs. **(B)** The Venn diagram shows overlap between genes with increased post-TES read density and *rdd* (*ros1-3 dml2-1 dml3-1*) hyper-methylated regions (relative to Col-0).

**Suppl Figure 6:**
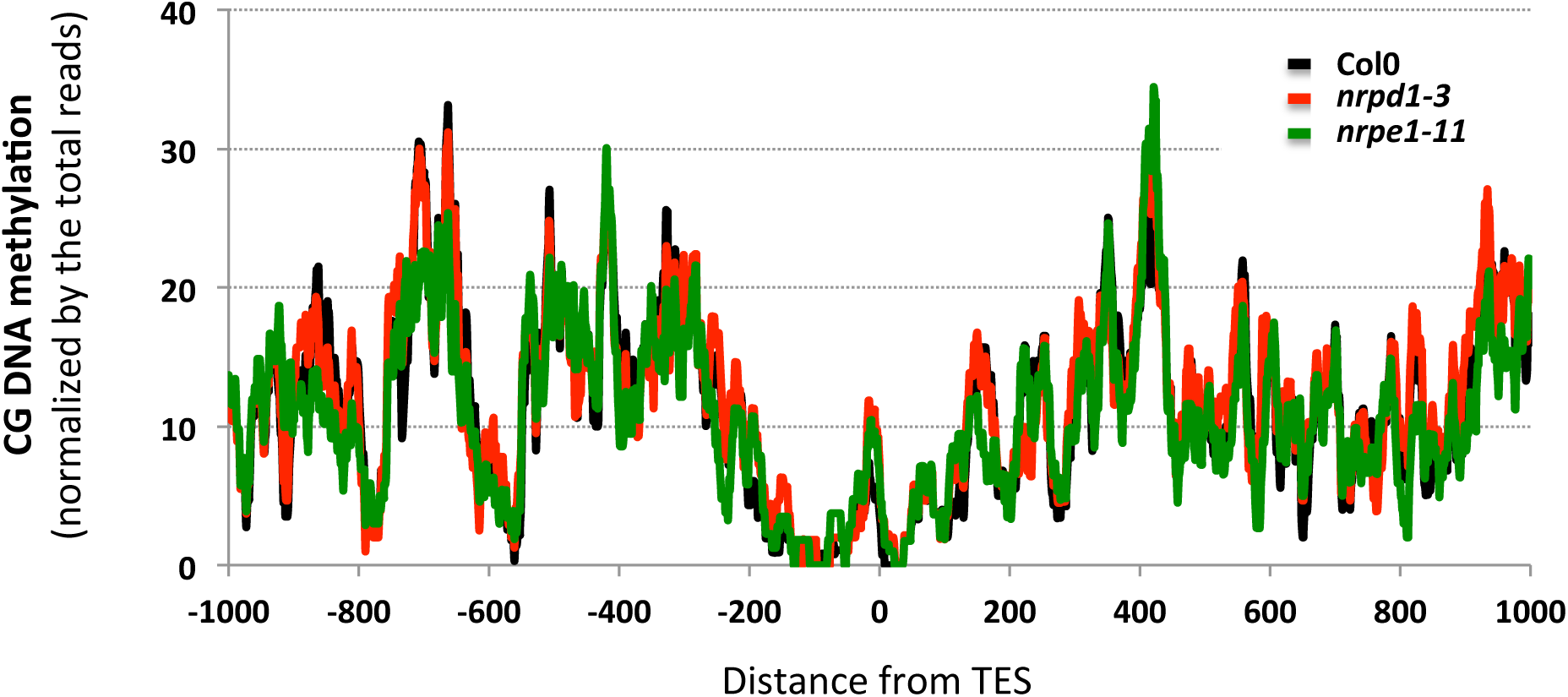
CG DNA methylation profile for overlapped set of 1312 genes with increased post-TES read density (by 2 fold or more) in mutant plants relative to wild-type Col-0 plants.

**Suppl Figure 7:**
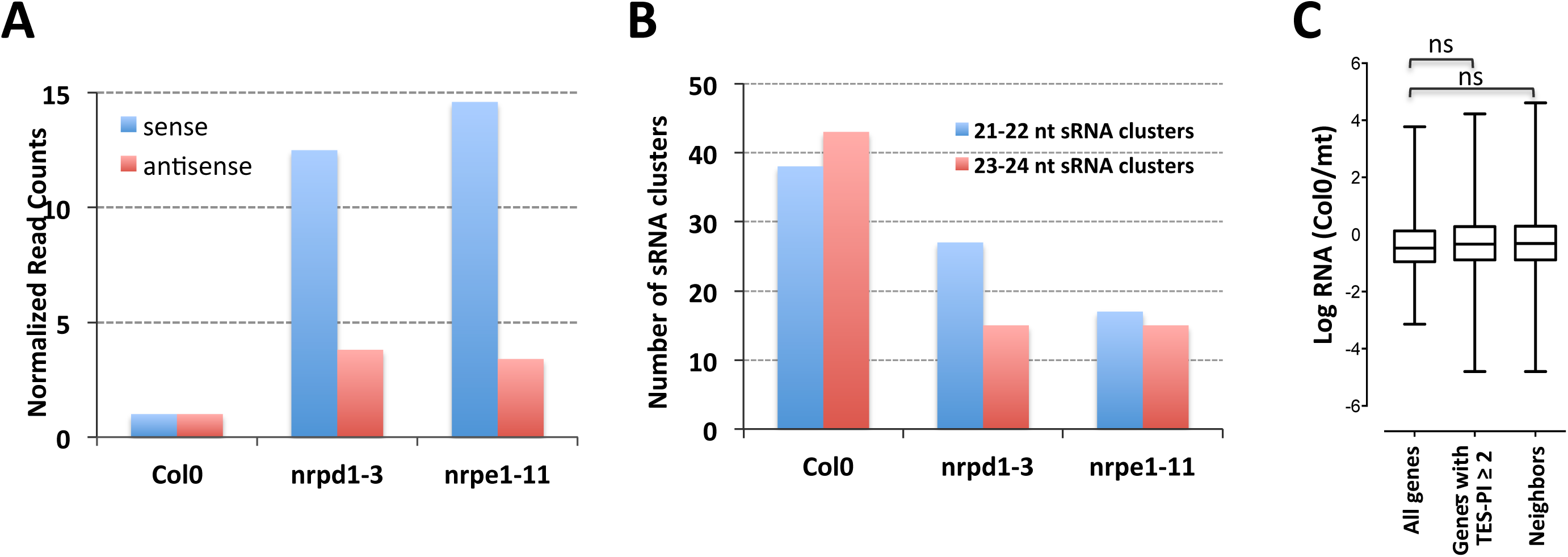
Increase in post-TES read density in *pol IV* or *pol V* mutants doesn’t affect transcript abundance for either gene in the convergent gene pairs. **(A)** The bar graph shows number of sense and antisense read counts detected within 1 kb regions surrounding TESs of genes with increased post-TES read density in Col-0, *pol IV* (*nrpd1-3*), and *pol V* (*nrpe1-11*) plants. **(B)** The bar graph compares number of 21-22 nt or 23-24 nt sRNA clusters identified within 1 kb regions surrounding TESs of genes with increased post-TES read density between Col-0, *pol IV* (*nrpd1-3*), and *pol V* (*nrpe1-11*) plants. **(C)** The plot shows Log_2_-transformed ratios of RNA read density (Col0/mt) for all protein-coding genes, genes with TES-TI (mt/Col0) ≥ 2, and their convergent neighbors. Significance was tested using ANOVA.

**Suppl Figure 8:**
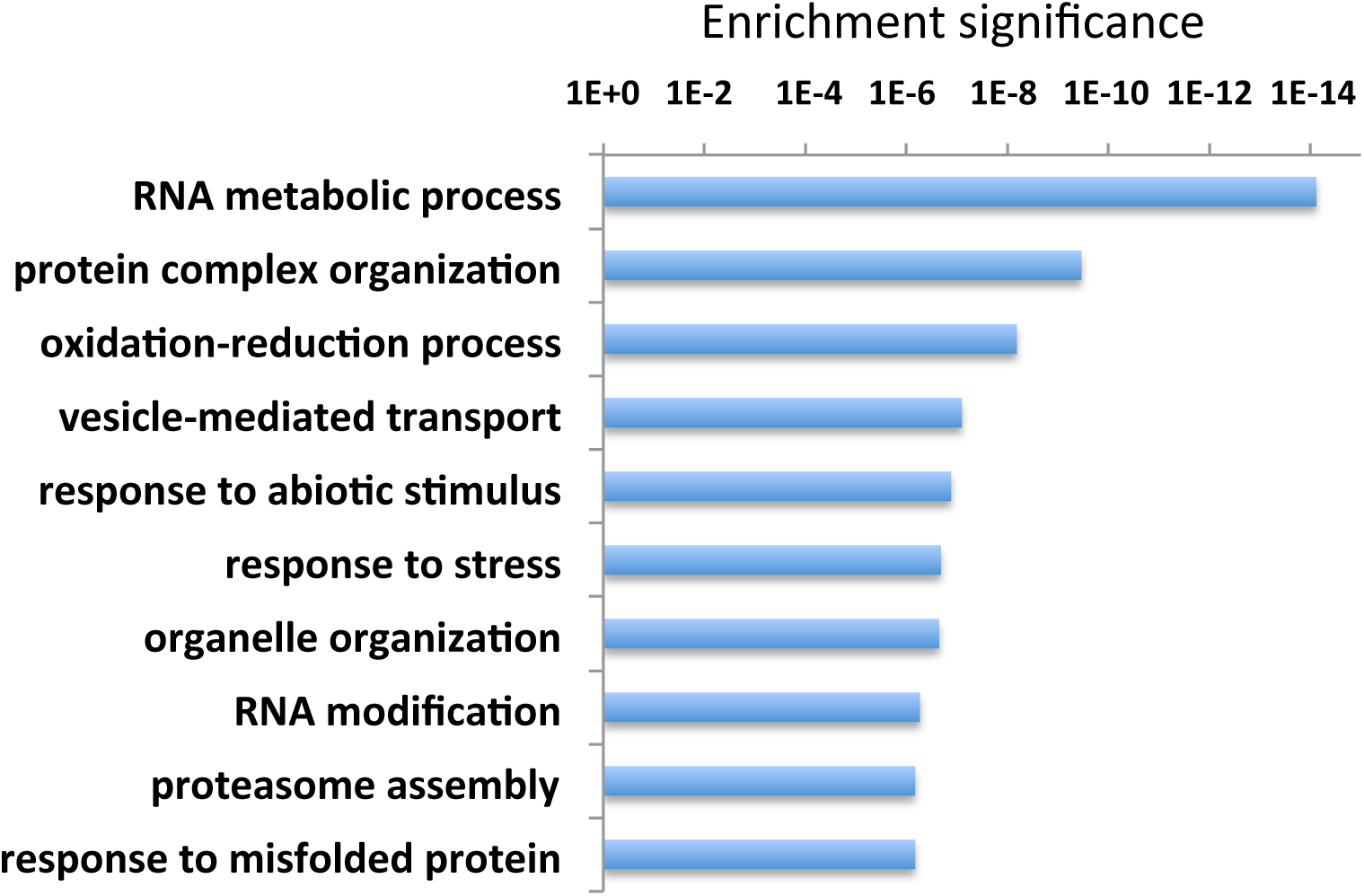
GO-enrichment analysis of transcripts with increased read density post-TES in *pol IV* or *pol V* mutant plants.

**Suppl Figure 9:**
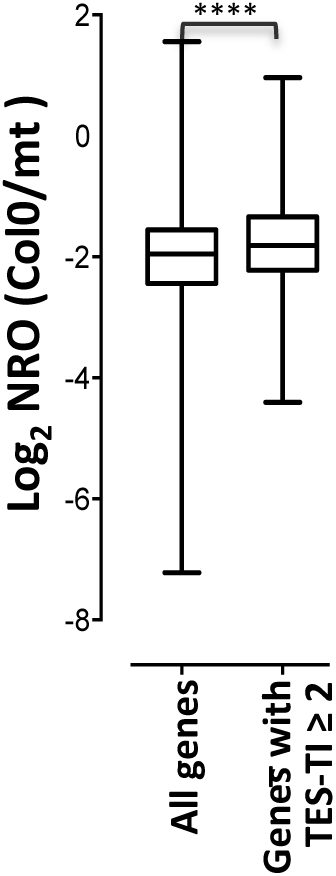
Transcripts with increased read density post-TES in *pol IV* or *pol V* mutant plants exhibit a significant increase in RNA synthesis in mutant plants. The plot shows Log_2_-transformed ratios of NRO read density (Col0/mt) for all protein-coding genes and genes with TES-TI (mt/Col0) ≥ 2. *P*-value (< 0.0001) was calculated using ANOVA.

**Suppl Figure 10:**
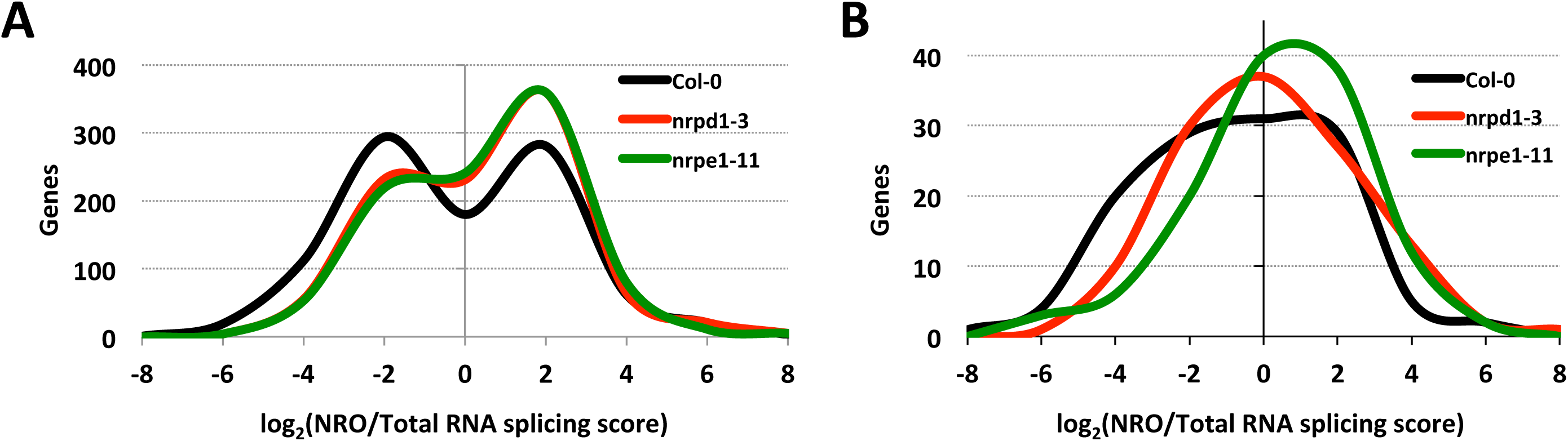
Splicing is less abundant in *pol IV* or *pol V* mutant plants. **(A)** The plot shows a distribution of the log_2_-transformed ratios of NRO to total RNA splicing scores. Splicing score was calculated in NRO and total RNA libraries as the ratio of the mean intronic read depth over the mean read depth in flanking exons for each intron-containing gene with RPKM of at least 4 (N = 973) to exclude bias from decreased intensity values toward gene ends. 1 indicates absence of splicing in the transcript and 0 indicates the transcript is fully spliced. **(B)** As in **(A)** but for transcripts with increased read density post-TES in *pol IV* or *pol V* mutants (N = 121).

